# Retrotransposons Promote *Dnmt3a*-Mutant Clonal Hematopoiesis Through Aging-Related Stromal Inflammation

**DOI:** 10.64898/2026.02.23.707588

**Authors:** Qiongdi Zhang, Wenhuo Hu, Joshua Theriot, Michael Lee, Sabrina Rinaldi, Yoon Jung Kim, Donghong Cai, Kathryn Dickerson, Jennifer Trowbridge, Nan Yan, Jian Xu, Sisi Zheng

**Author notes:** **Corresponding Author**: Sisi Zheng, MD, 5323 Harry Hines Blvd, Dallas, Texas, 75390. **Data Sharing Statement**: RNA-sequencing data have been deposited in the Gene Expression Omnibus (accession number GSE325131).

## Abstract

Clonal hematopoiesis (CH) is an age-related phenomenon driven by the expansion of mutant hematopoietic stem cell (HSC) clones, most commonly harboring mutations in *DNMT3A*. While inflammation is known to promote CH, the upstream triggers of this inflammatory state remain unclear. We show that aging selectively upregulates retrotransposons in the non-hematopoietic cell compartment of the murine bone marrow, particularly in mesenchymal stromal cells. Using *in vivo* competitive transplant models, we demonstrate that retrotransposon-induced inflammation cell-extrinsically promotes *Dnmt3a*-mutant HSC expansion. This competitive advantage arises from mutant HSC resistance to inflammation-driven cell cycle perturbations. Mechanistically, we show that retrotransposon activation induces type I interferon signaling via viral mimicry, such that stromal knockdown of either *Irf3* or *Sting* abrogates the competitive advantage of *Dnmt3a*-mutant HSCs. Our findings establish stromal-selective retrotransposon reactivation as a previously unrecognized, non-cell-autonomous source of inflammation that contributes to age-associated CH.

**Key Points:**

- Retrotransposons remodel the aging niche through viral mimicry pathways to promote *Dnmt3a*-mutant clonal hematopoiesis.
- Retrotransposon-mediated inflammation impairs wild-type HSC fitness, while *Dnmt3a*-mutant HSCs resist inflammatory changes.

## Introduction

Clonal hematopoiesis (CH) is characterized by the clonal expansion of hematopoietic stem cell (HSC) clones bearing fitness-conferring mutations and leads to an increased risk for progression to acute myeloid leukemia (AML) and the myelodysplastic syndromes (MDS).^1,2^ More than half of CH cases harbor mutations in DNA methyltransferase 3A (*DNMT3A*), most notably the hotspot mutation *DNMT3A*^*R882H*^. CH is closely associated with aging, and mounting evidence implicate HSC-intrinsic and HSC-extrinsic mechanisms of inflammation as critical promoters of CH.^3-9^ For example, TNFα and IFNγ can drive *Dnmt3a*-mutant CH, and mutant HSCs can also generate pro-inflammatory mature progeny, reinforcing a CH-promoting positive feedback loop.^5,10-13^ Nonetheless, the cellular and molecular origins of the aging-associated inflammation that promote CH remain poorly understood.

An underexplored contributor to sterile inflammation in aging tissues are retrotransposons, which include families of long interspersed nuclear elements (LINEs), short interspersed nuclear elements (SINEs), and endogenous retroviruses (ERVs). Retrotransposons are genomic elements capable of reverse transcription, enabling their sequences to be ‘copy-and-pasted’ across the genome. These mobile elements are normally heavily silenced in somatic cells.^14^ However, aging can lead to their reactivation, thereby triggering a viral mimicry response (e.g. cGAS-STING and RIG-I-like receptor pathways) and inducing type I interferons (IFN-I), including IFNα/β.^15-17^ Recent work has linked retrotransposon-induced inflammation with cancer therapy responsiveness and stress-induced erythropoiesis.^18-21^ The expression of retrotransposons has also been shown to be tightly and intrinsically regulated in AML cells.^22,23^ However, whether retrotransposons can drive CH by promoting aging-associated inflammation remains unknown.

Here, we investigate whether and how retrotransposons within the aged bone marrow environment contribute to the selective expansion of *Dnmt3a*-mutant HSCs. We find that during aging, mesenchymal stromal cells (MSCs) selectively upregulate retrotransposons and IFN-I. Using *in vivo* and *ex vivo* models, we show that retrotransposons non-cell-autonomously promote the competitive fitness of *Dnmt3a*-mutant HSCs through a viral mimicry response. These findings identify a novel extrinsic driver of CH, linking age-associated retrotransposons in the bone marrow stroma to the selective expansion of mutant HSCs.

## Methods

### Mice

C57BL/6J, *Ptprc*^*em6Lutzy*^/J CD45.1, B6.Cg-*Commd10*^*Tg(Vav1-icre)A2Kio*^/J, and B6(Cg)-*Ifnar1*^*tm1*.*1Ees*^/J mice were obtained from The Jackson Laboratory. *Mpp8*^*-/-*^ mice were provided by Dr. Jian Xu.^23^ *Dnmt3a*^*fl-R878H/fl-R878H*^ mice (provided by Dr. Jennifer Trowbridge)^24^ were crossed with *Vav-iCre* mice and CD45.1 to generate CD45.1/CD45.2 *Vav-iCre*; *Dnmt3a*^*fl-R878H/+*^ progeny, or crossed with *Ifnar1*^*fl/fl*^ mice to obtain *Vav-iCre*;*Dnmt3a*^*fl-R878H/+;*^ *Ifnar1*^*fl/fl*^ and *Vav-iCre*;*Ifnar1*^*fl/fl*^. All animal procedures were approved by the Institutional Animal Care and Use Committee at UTSW and carried out in accordance with institutional guidelines.

### MSC and EC isolation

Pelvis and long bones were dissected and flushed with HBSS with Ca^2+^/Mg^2+^. Marrow plugs were digested in 2.5mg/mL Collagenase A, 4mg/mL Dispase II, and 0.01mg/mL DNase I at 37°C for 20 minutes with shaking. The suspension was filtered (70µm strainer) and deactivated with HBSS with 2% FBS and 2mM EDTA. Cells were washed and stained for FACS sorting (see Table S1 for antibodies).

### *In vivo* assays

Recipient mice were conditioned with IP busulfan (20mg/kg/day, day -5, -4, -3). WBM cells from CD45.1 mice and CD45.1/CD45.2; *Vav-iCre*; *Dnmt3a*^*fl-R878H/+*^ mice were mixed at a 5:1 ratio and retro-orbitally injected into recipient *Mpp8*^*-/-*^ or *Mpp8*^*+/+*^ mice on day 0. Peripheral blood was analyzed monthly by CBC machine (ProCyte Dx) and by FACS (BD FACSymphony). At five months post-transplant, mice were euthanized for BM cell harvest. Absolute HSC quantification was performed using bone marrow plugs flushed directly from long bones and subjected to ACK lysis. Total nucleated cell number was obtained using Nexcelom Cellometer Auto 2000 cell counter and multiplied by HSC frequency obtained by flow analysis to calculate absolute HSC count. CFU assays and serial replating assays were performed using pre-aliquoted 3mL vials of Methocult M3434 in a 6-well plate. BM cells were isolated and stained (see Table S1 for antibodies).

### *Ex vivo* assays

LT-HSCs (CD150^+^CD34^−^LSK) from WT (CD45.2) and *Dnmt3a*-mutant (CD45.1/CD45.2) mice were isolated by flow cytometry and mixed at a 2:1 ratio. 1200 mixed LT-HSCs were plated onto a confluent layer of MS5 stromal cells in 24-well plates. Co-cultures were maintained in StemSpan SFEM II supplemented with SCF (50ng/mL) for 11 days. Half-media exchange was performed on Day 5 and Day 8. On Day 11, non-adherent cells were harvested, stained, and analyzed by flow cytometry. Donor chimerism was calculated as the percentage of mutant (CD45.1/CD45.2) cells among total hematopoietic cells. Antibody details are provided in Table S1. For overexpression studies, retrotransposon consensus sequences for Lx8, B1, and MERVL were identified from RNA sequencing data and cloned into the pCAGGS-Puro-mCherry lentiviral vector. An empty vector (EV) was used as the control. MS5 stromal cells were transiently transfected with these constructs using Lipofectamine 3000. For the shRNA knockdown studies, shRNA sequences targeting *Irf3* and *Sting* were designed based on previously published sequences and shLuciferase served as control.^25^ The shRNA oligonucleotides were synthesized, annealed, and cloned into the doxycycline-inducible pTRIPZ-TagBFP-miR-E-hPGK-TetOn3G-2A-PuroR lentiviral vector.

## Results

### Aging activates retrotransposons and inflammation in the bone marrow environment

We first sought to identify cellular and molecular sources of bone marrow inflammation during aging that could act as putative drivers of CH. To distinguish cell-intrinsic from cell-extrinsic contributors to CH, we harvested hematopoietic and non-hematopoietic bone marrow cells from two-month-old young mice and 24-month-old aged mice for bulk RNA sequencing (Figure 1A, Figure S1A-B). Within the non-hematopoietic compartment, we separated the two major cell types, mesenchymal stromal cells (MSCs) and endothelial cells (ECs); and within the hematopoietic compartment, we enriched for cKit^+^ hematopoietic stem and progenitor cells (HSPCs) (Figure 1B). We found that aging transcriptionally altered MSCs to a much greater extent than ECs and HSPCs (Figure S1C-D). Moreover, pathways enriched with aging in MSCs and ECs were dominated by innate immune signaling programs, especially TNFα and IFNα responses (Figure 1C-D). In contrast, aged HSPCs showed no significant enrichment for inflammatory pathways, suggesting aging-associated inflammation primarily stems from constituents of the bone marrow environment (Figure 1C).

**Figure 1.**
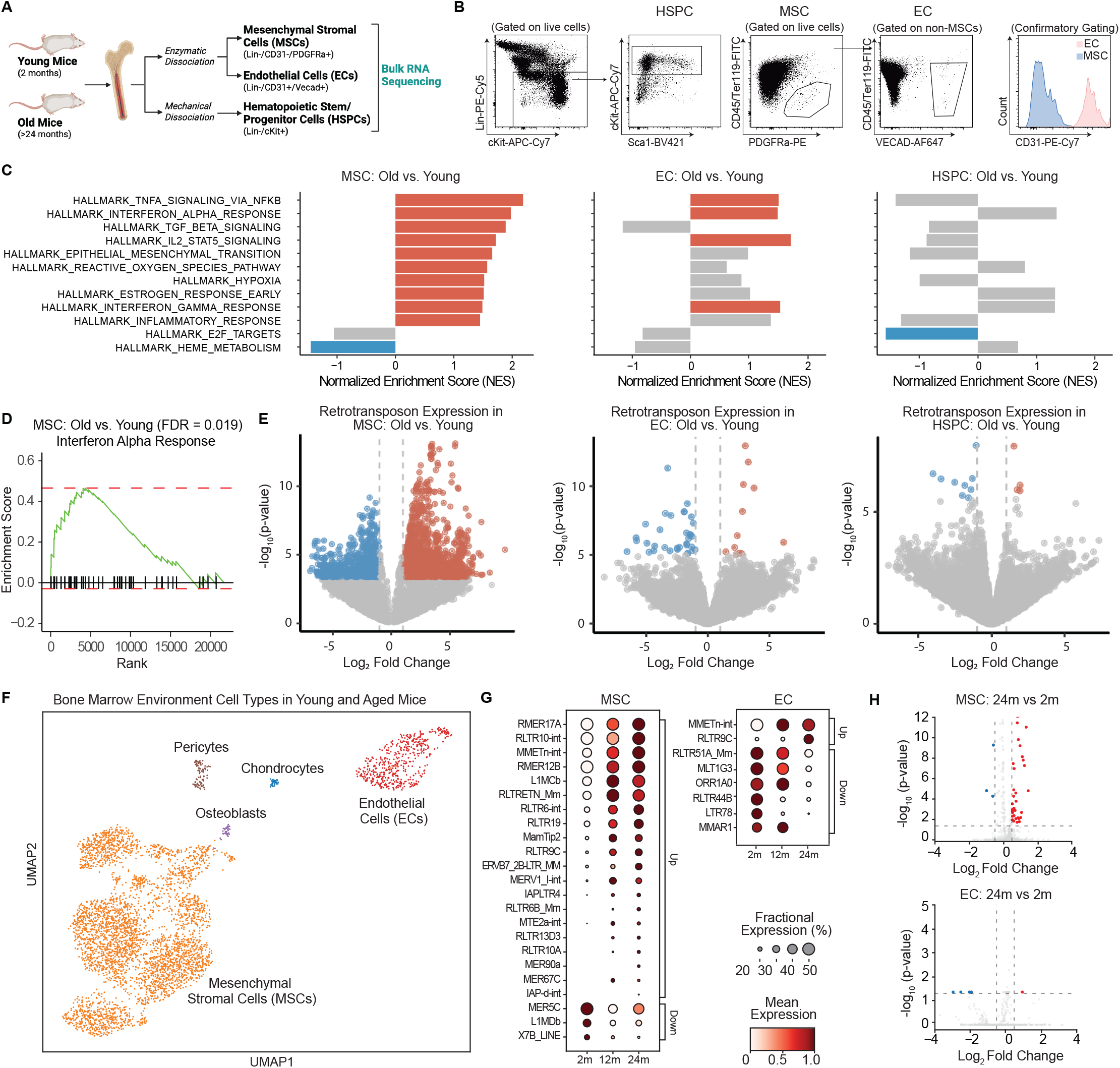
Aging activates retrotransposons and inflammation in the bone marrow environment. **(A)** Schematic of cell processing for RNA sequencing of old and young murine bone marrow cells (n=4-5). Image created using BioRender. (**B)** Flow cytometry gating strategy for sorting HSPCs, MSCs, and ECs. CD31 was used to confirm separation of MSC and EC populations. HSPC, hematopoietic stem and progenitor cells; MSC, mesenchymal stromal cells; EC, endothelial cells. **(C)** Gene set enrichment analyses comparing bone marrow cell types from old versus young mice. Red and blue bars indicate significant up- and down-regulation, respectively. Grey bars indicate non-significant changes. Significance threshold set at FDR < 0.1. **(D)** Positive enrichment for genes related to the interferon alpha response in aged MSCs. **(E)** Volcano plot showing differentially expressed retrotransposon loci in old versus young mice across bone marrow cell types. **(F)** UMAP clustering analysis of a public scRNA-seq dataset (GEO: GSE299733) identifying 5 bone marrow environment cell types in young (2 month), middle age (12 month), and old (24 month) mice. **(G)** Dot plots comparing fractional and mean expression of retrotransposons in 2-month-old versus 24-month-old MSCs and ECs. **(H)** Volcano plot depicting differentially expressed retrotransposon genes from (G) with assigned significance values.

Given that retrotransposons have been implicated in aging-related IFN-I signaling, we next examined whether retrotransposon upregulation was correlated with the observed IFNα induction in the bone marrow environment.^15,16^ Indeed, aging preferentially upregulated retrotransposons in MSCs over ECs, and HSPCs exhibited only minor changes in retrotransposon expression (Figure 1E). To further validate these findings, we reanalyzed publicly available single-cell-RNA-sequencing data from the bone marrow environment of young (two-month-old), middle-aged (12-month-old), and old (24-month-old) mice, which previously identified increased IFN signatures in the aged MSC populations (Figure 1F).^26^ Our analyses confirmed that MSCs selectively upregulate retrotransposons with aging. Notably, a substantial fraction of retrotransposons already exhibited increased expression by middle age, with levels continuing to trend upward in old age (Figure 1G). Additional sub-clustering of MSCs revealed heterogeneity in retrotransposon upregulation across stromal subpopulations (Figure S1E-G). In contrast, aged ECs, osteoblasts, and pericytes showed minimal changes in retrotransposon expression when examined at both the 12-month and 24-month timepoints (Figure 1H, Figure S1H). Together, these bulk and single-cell RNA sequencing data demonstrate that retrotransposon upregulation in the aging bone marrow is largely restricted to non-hematopoietic cells, particularly MSCs, and is tightly associated with a robust interferon response.

### Aged stromal cells preferentially upregulate retrotransposons from intronic regions

To determine how retrotransposons interact with host gene activities during aging, we assessed their genomic distribution. Intriguingly, nearly 80% of retrotransposons upregulated in aged MSCs were located within intronic regions, whereas fewer than 30% of downregulated retrotransposons mapped to introns and were more common in intergenic regions (Figure S2A). Consistent with this pattern, aging selectively increased intron-derived retrotransposon expression in MSCs, a shift not observed in ECs or HSPCs (Figure S2B). Because intron retention is a recognized hallmark of aging-associated splicing dysfunction, we also examined host splicing patterns and found widespread accumulation of intronic sequences in aged MSCs (Figure S2C).^27,28^ To validate our RNA sequencing results, we performed RT-qPCR on an ERV-K family retrotransposon, *MMETn-int*, located within an intron of the interferon-stimulated gene *Gvin1*. Aging significantly increased intronic *MMETn-int* expression through intron retention in MSCs but not in unfractionated whole bone marrow (Figure S2D-E), supporting a MSC-specific splicing effect. We then investigated potential mechanisms underlying this phenomenon and identified a subset of splicing factors that were dysregulated with aging in MSCs. In contrast, HSPCs exhibited higher baseline expression of splicing factors that remained stable with aging, suggesting that MSCs may be intrinsically more vulnerable to splicing perturbations (Figure S2F). Our findings support a cell-type-specific link between aging-associated splicing alterations and intron-derived retrotransposon activation in MSCs.

### Retrotransposons non-cell-autonomously promote *Dnmt3a*-mutant HSC expansion

We next investigated whether retrotransposons can influence the growth dynamics of CH in a non-cell-autonomous manner. To directly test whether retrotransposons in the bone marrow can induce an inflammatory response, we used a genetically engineered mouse model with constitutive loss of *Mpp8*, a member of the HUSH complex which epigenetically mediates retrotransposon silencing. Bulk RNA sequencing of unfractionated bone marrow cells from *Mpp8*^*-/-*^ mice revealed broad upregulation of retrotransposons accompanied by a strong innate immune response and interferon signature in both the whole bone marrow and in MSCs (Figure 2A-C, Figure S3A-B).^23^

**Figure 2.**
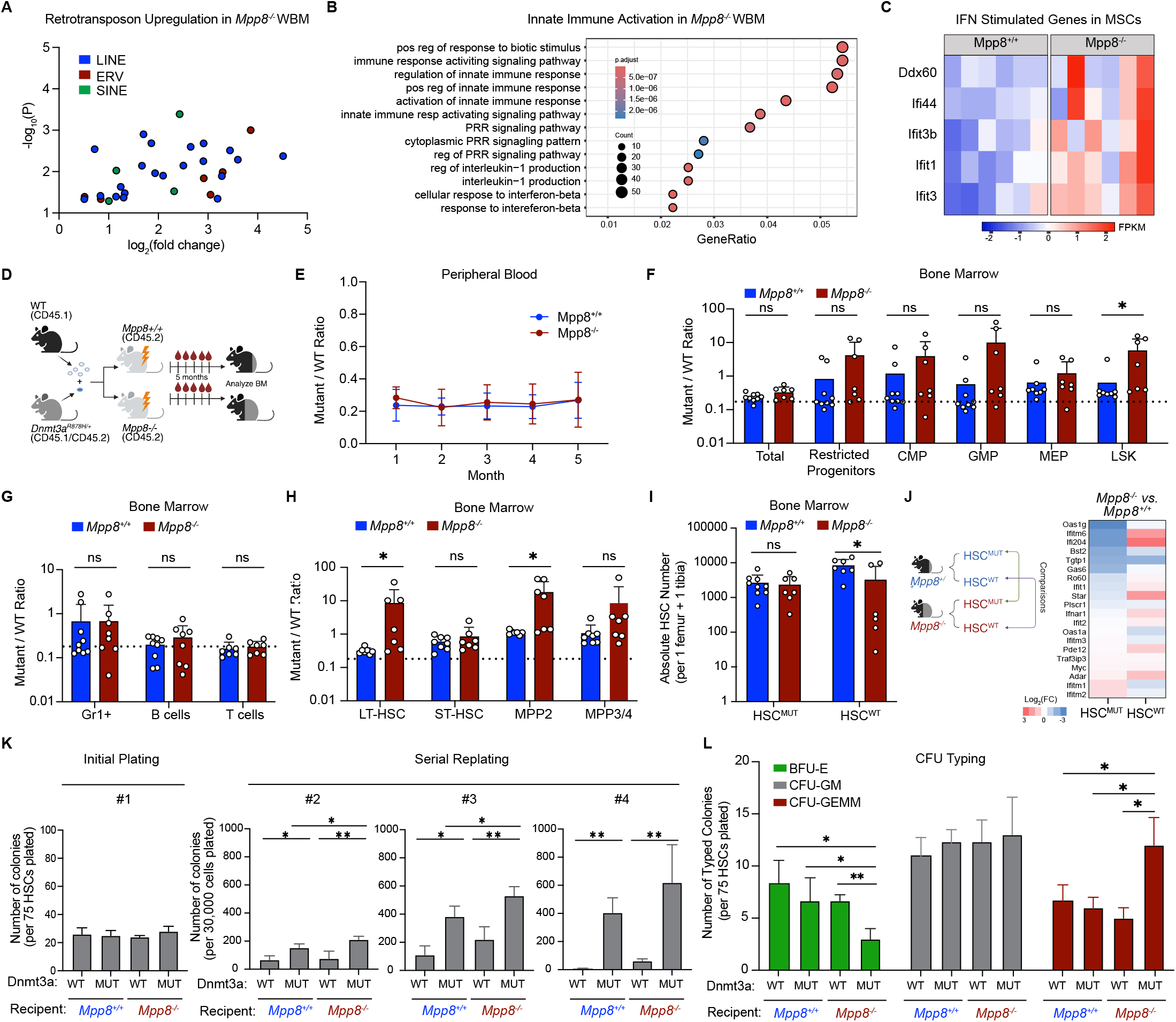
Retrotransposons non-cell-autonomously promote *Dnmt3a*-mutant HSC expansion. **(A)** Upregulated retrotransposons (color-coded by retrotransposon family) in the WBM of *Mpp8*^-/-^ versus *Mpp8*^+/+^ mice (n=4-6). WBM, whole bone marrow. **(B)** Gene set enrichment analysis comparing *Mpp8*^-/-^ WBM versus *Mpp8*^+/+^ WBM. **(C)** Gene expression (FPKM) of type I interferon-related genes in *Mpp8*^-/-^ MSCs versus *Mpp8*^+/+^ MSCs. **(D)** Schematic of competitive bone marrow transplant in which WBM from *Dnmt3a*-mutant and WT mice are mixed at a 1:5 ratio and transplanted into either *Mpp8*^-/-^ and *Mpp8*^+/+^ recipients conditioned with busulfan (n=6-8). Images created using BioRender. **(E)** Monthly peripheral blood analysis of *Dnmt3a*-mutant/WT total cell ratio between *Mpp8*^-/-^ and *Mpp8*^+/+^ recipients by flow cytometry (n=6-8). **(F-H)** Endpoint bone marrow analysis by flow cytometry of *Dnmt3a*-mutant/WT ratio between *Mpp8*^-/-^ and *Mpp8*^+/+^ recipients across total cell, stem/progenitor populations (F); mature cell compartments (G); and HSC/MPP compartments (H). Statistical significance was determined by multiple t-tests (n=6-8). CMP, common myeloid progenitor; GMP, granulocyte-macrophage progenitor; MEP, megakaryocyte-erythroid progenitor; LT-HSC, long-term HSC; ST-HSC, short-term HSC; MPP, multipotent progenitor. **(I)** Quantification of absolute numbers of *Dnmt3a*-mutant and WT HSCs from *Mpp8*^-/-^ and *Mpp8*^+/+^ recipient groups at endpoint analysis. Statistical significance was determined by multiple t-tests (n=6-8). **(J)** *Dnmt3a*-mutant (HSC^MUT^) and WT HSCs (HSC^WT^) were collected at the end of the competitive bone marrow transplant (described in Figure 2D) for RNA-seq. Comparisons were made as shown (left) to generate heatmaps (right) of differentially expressed interferon-stimulated genes. Fold change in gene expression is a non-statistical comparison of pooled HSCs for low-input RNA-seq across all biological samples to permit adequate depth of analysis. **(K)** 75 donor-derived *Dnmt3a*-mutant HSCs or WT HSCs from *Mpp8*^*-/-*^ or *Mpp8*^*+/+*^ recipient groups were purified at the end of the competitive bone marrow transplant experiment (as described in Figure 2D) and plated into methylcellulose in 6-well plates. Total colonies were counted at the end of 10 days (n=3). 30,000 cells from the prior plating were then serially replated into fresh methylcellulose wells and total colonies were counted every 10 days. **(L)** Initial plating (in Figure 2K) was scored for CFU-GEMM, CFU-GM, and BFU-E colonies (n=3).

Next, we tested whether retrotransposon-associated inflammation in the bone marrow environment of *Mpp8*^*-/-*^ mice could favor the selective expansion of mutant HSCs *in vivo*. To this end, we utilized a well-established mouse model of CH harboring a heterozygous *Dnmt3a*^*R878H*^ mutation^24^, homologous to human *DNMT3A*^*R882H/+*^.^29^ We performed competitive transplants by mixing CD45.1/CD45.2 *Vav-iCre*;*Dnmt3a*^*fl-R878H/+*^ (hereafter referred to as *Dnmt3a*-mutant) and CD45.1 wild-type (WT) bone marrow cells at a 1:5 ratio and transplanting these cells into busulfan-conditioned CD45.2 *Mpp8*^*-/-*^ or *Mpp8*^*+/+*^ recipients (Figure 2D). In control experiments, we confirmed that busulfan itself did not lead to an IFN-I marrow response, and we optimized busulfan dosing to use the lowest dose that allowed for adequate engraftment of HSCs to minimize damage to the bone marrow stroma (Figure S3C-E).

Serial peripheral blood analysis over five months showed no difference in *Dnmt3a*-mutant (CD45.1/CD45.2) versus WT (CD45.1) chimerism between the *Mpp8*^*-/-*^ or *Mpp8*^*+/+*^ recipient groups in total cells or in T cells, B cells and myeloid cells (Figure 2E, Figure S3F). Peripheral blood counts were also comparable between groups (Figure S3G). Strikingly, terminal bone marrow evaluation at five months revealed a selective outgrowth of *Dnmt3a*-mutant cells in *Mpp8*^*-/-*^ recipients as compared to *Mpp8*^*+/+*^ recipients in the Lineage^-^Sca-1^+^cKit^+^ (LSK) compartment, a population that is highly enriched for HSCs (Figure 2F, gating strategy described in Figure S3H). This selective outgrowth of mutant cells was not observed in the unfractionated bone marrow cells, restricted progenitor populations, or mature cell populations, consistent with the peripheral blood (Figure 2F-G). The restriction of mutant cell outgrowth within the LSK cells reflects known differentiation defects associated with *Dnmt3a* loss.^30-32^ Within the LSK cells, we also observed selective outgrowth of *Dnmt3a*-mutant cells in the long-term-HSC and multipotent progenitor 2 (MPP2) populations, but no such selective outgrowth in short-term-HSC or MPP3/4 populations, indicating that retrotransposons may promote the selective expansion of specific HSC and MPP subsets (Figure 2H). To control for potential confounding effects of CD45 alleles or non-specific *Vav-iCre* activation, we performed parallel transplants competing a mixture of *Vav-iCre* (CD45.1/CD45.2) and WT (CD45.1) bone marrow cells. In this setting, we observed no selective outgrowth of *Vav-iCre* cells between the *Mpp8*^*-/-*^ or *Mpp8*^*+/+*^ recipients, confirming the specificity of the effect to *Dnmt3a*-mutant cells (Figure S3I-J).

Despite the selective outgrowth of *Dnmt3a*-mutant HSCs over WT HSCs in *Mpp8*^*-/-*^ but not *Mpp8*^*+/+*^ recipients, there was no difference in the absolute number of *Dnmt3a*-mutant HSCs between the two recipient groups in the terminal analyses. Conversely, the absolute number of WT HSCs was reduced in the *Mpp8*^*-/-*^ compared to *Mpp8*^*+/+*^ recipients (Figure 2I). This suggests that retrotransposons in the bone marrow stroma enhance the competitive advantage of *Dnmt3a*-mutant HSCs by impairing WT HSC fitness. To further explore this possibility, we separately isolated engrafted WT (CD45.1) and *Dnmt3a*-mutant (CD45.1/CD45.2) HSCs from the *Mpp8*^*-/-*^ or *Mpp8*^*+/+*^ recipients at the end of the five-month competitive transplant experiment and performed bulk RNA sequencing. Indeed, WT HSCs from *Mpp8*^*-/-*^ recipients exhibited increased interferon-stimulated gene (ISG) expression compared to WT HSCs from *Mpp8*^*+/+*^ recipients, while much less ISG activation was observed in *Dnmt3a-*mutant HSCs (Figure 2J). These results suggest that WT HSCs are reprogrammed by retrotransposon-induced inflammatory stress in the aged bone marrow environment, whereas *Dnmt3a*-mutant HSCs are comparatively resistant.

To assess HSC functional capacity, we performed methylcellulose-based colony-forming assays using the HSCs isolated at the endpoint of the competitive transplant experiments. As expected, *Dnmt3a*-mutant HSCs generated more colonies than WT HSCs upon serial replating.^32^ Moreover, *Dnmt3a*-mutant HSCs transplanted into *Mpp8*^*-/-*^ recipients formed more CFU-GEMM colonies compared to *Dnmt3a*-mutant HSCs transplanted into *Mpp8*^*+/+*^ recipients, and they also exhibited more robust colony-forming ability in serial replating assays (Figure 2K-L). Hence, *Dnmt3a*-mutant HSCs maintain greater clonogenic potential under inflammatory conditions induced by retrotransposons. These findings demonstrate that retrotransposon upregulation in the bone marrow environment not only selectively depletes WT HSCs but it also promotes the functional potential of *Dnmt3a*-mutant HSCs, implicating retrotransposons as an age-associated extrinsic factor in the evolution of CH.

### *Dnmt3a*-mutant HSCs resist cell cycle changes driven by retrotransposon-induced inflammatory stress

To further explore the causal link between retrotransposons and CH progression, we asked whether ectopic overexpression of an aging-related retrotransposon in bone marrow stromal cells is sufficient to promote the selective outgrowth of *Dnmt3a*-mutant HSCs. To test this, we devised a co-culture assay in which HSCs could be expanded *ex vivo* for at least eleven days on a monolayer of MS5 stromal cells overexpressing a single retrotransposon. From our previous RNA sequencing analysis of young and old mouse bone marrow (Figure 1E), we identified Lx8, a murine LINE-1 family retrotransposon, to be highly upregulated in MSCs during aging (Figure S4A). Thus, we cloned the sequence of Lx8 and overexpressed this element in MS5 mouse stromal cells. After verifying robust overexpression of Lx8 in the MS5 cells (Figure S4C), we purified *Dnmt3a*-mutant and WT HSCs (CD150^+^CD34^-^LSKs) from mice and seeded them in a 1:2 ratio onto a monolayer of MS5 cells expressing either empty vector (EV) or Lx8 (hereafter referred to as MS5-EV and MS5-Lx8). After eleven days, we assessed the competitive growth of *Dnmt3a*-mutant and WT cells by flow cytometry (Figure 3A). While Lx8 overexpression in MS5 stromal cells did not alter the relative abundance of *Dnmt3a*-mutant cells in the total or restricted progenitor compartments, it significantly enhanced the fitness advantage of *Dnmt3a*-mutant cells in the LSK compartment (Figure 3B-C). Notably, the absolute number of *Dnmt3a*-mutant LSK cells did not change between co-culture with MS5-EV or MS5-Lx8 cells, whereas the absolute number of WT LSK cells was reduced in the MS5-Lx8 co-culture group (Figure 3D). These experiments recapitulate the results of our *in vivo* experiments transplanting *Dnmt3a*-mutant cells into *Mpp8*^*-/-*^ recipients (in Figure 2I). Moreover, they affirm that WT cells are more sensitive than *Dnmt3a*-mutant cells to the cell-extrinsic effects of retrotransposons, resulting in selective impairment of WT LSK expansion. Thus, our data support a model in which retrotransposons promote the competitive advantage of *Dnmt3a*-mutant clones primarily by suppressing WT HSC fitness rather than by enhancing mutant cell proliferation.

**Figure 3.**
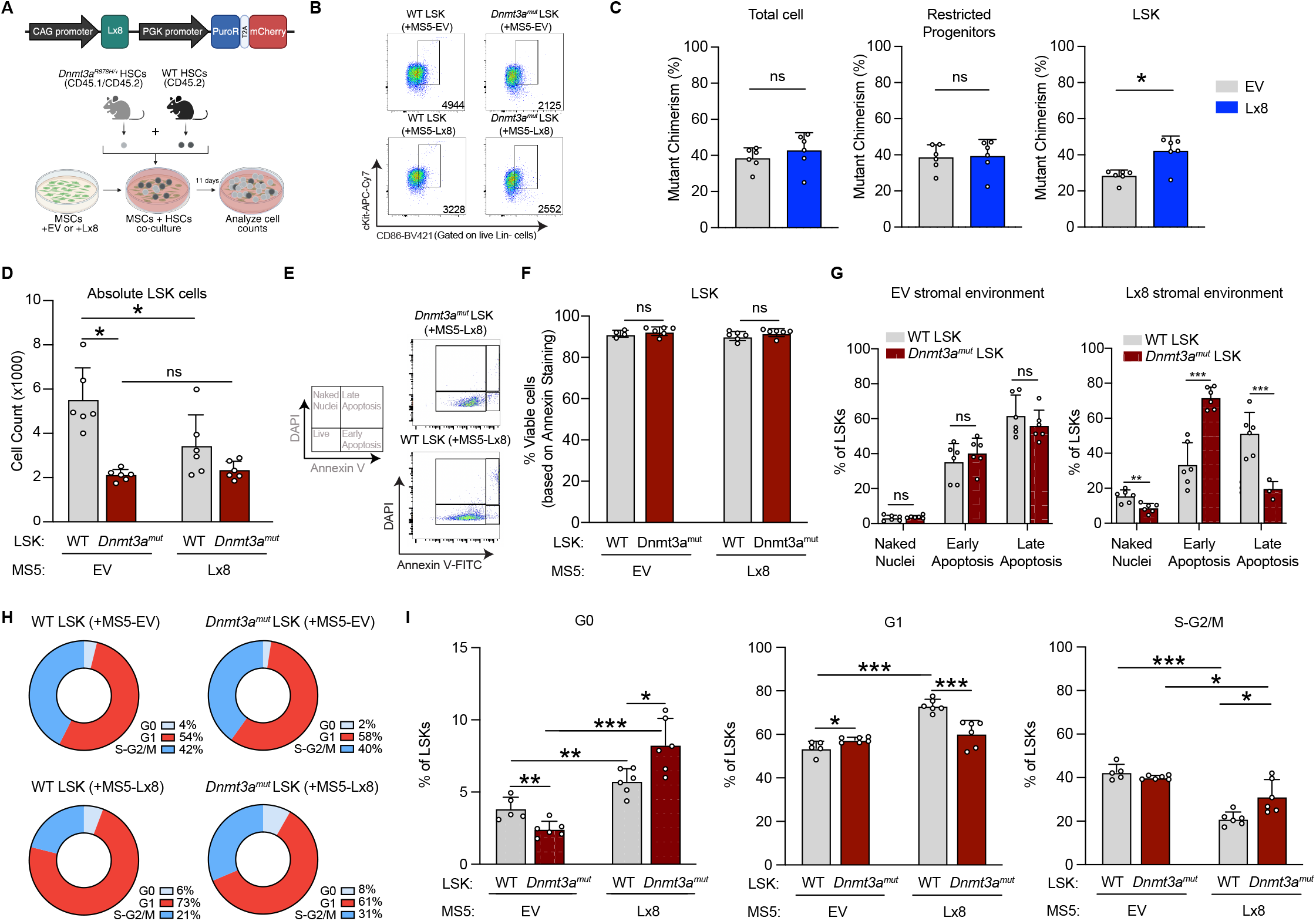
*Dnmt3a*-mutant HSCs resist cell cycle changes driven by retrotransposon-induced inflammatory stress. **(A)** Schematic of Lx8 overexpression vector (top) and *ex vivo* competitive co-culture model (below). LT-HSCs (SLAM^+^CD34^−^ LSKs) were sorted from WT CD45.2 mice and *Vav-iCre Dnmt3a*^*fl-R878H/+*^ CD45.1/CD45.2 mice. WT (CD45.2) and *Dnmt3a*-mutant (CD45.1/CD45.2) LT-HSCs were mixed at a 2:1 ratio and co-cultured with MS5 cells transfected with an empty vector or the Lx8 construct. After 11 days of co-culture, HSCs were analyzed by flow cytometry. **(B)** Representative flow cytometry gating to determine LSK counts from co-culture experiment described in (A). CD86 was used in place of Sca-1 staining for LSK cells due to IFNα-induced Sca1 upregulation. Absolute cell numbers are shown in bottom right of each plot. **(C)** Flow cytometry analysis of mutant chimerism in total cells, restricted progenitors (LK), and LSKs after *ex vivo* co-culture (n=5-6). Mutant chimerism was calculated as the ratio of *Dnmt3a*-mutant CD45.1/CD45.2 cells to total CD45 cells within the defined compartment. Statistical significance was determined by multiple t-tests. **(D)** Quantification of absolute LSK numbers after mixed HSCs were co-cultured with MS5-EV and MS5-Lx8 (n=5-6). Statistical significance was determined by multiple t-tests. **(E)** Flow cytometry gating scheme (left) and representative flow plots (right) for determining early and late apoptosis frequency in LSK cells. Analysis was conducted by flow cytometry using DAPI and Annexin-V. **(F)** Overall cell viability in WT and mutant LSK populations co-cultured with MS5-EV and MS5-Lx8 **(G)** Cell frequency of early/late apoptosis and naked nuclei in WT and mutant LSK populations co-cultured with MS5-EV (left) and MS5-Lx8 (right). **(H)** Pie charts depicting the cell cycle distribution of *Dnmt3a*-mutant and WT LSKs co-cultured with MS5-EV and MS5-Lx8. Analysis conducted by flow cytometry using DNA content and nuclear Ki-67 staining. **(I)** Bar chart quantification of the pie chart data in (H), comparing cell cycle phase frequencies of LSKs co-cultured with MS5-EV and MS5-Lx8 (n=5-6). **(J)** Pie charts depicting the cell cycle distribution of *Dnmt3a*-mutant and WT HSCs following 7 days of *in vivo* IFNα (4×10^5^ IU/kg/day) or vehicle treatment. **(K)** Bar chart quantification of the pie chart data in (J), comparing cell cycle phase frequencies of *Dnmt3a*-mutant and WT HSCs following 7 days of *in vivo* IFNα (4×10^5^ IU/kg/day) or vehicle treatment. (n=5-6). Statistical significance was determined by multiple t-tests.

To understand whether the impaired fitness of WT LSKs in retrotransposon-activated environments results from changes in cell death or cell growth, we assessed apoptosis and cell cycle status following *ex vivo* co-culture. While MS5-Lx8 exposure did not change the overall frequency of apoptosis in the WT or *Dnmt3a*-mutant LSKs, WT LSKs were biased toward late apoptotic states and *Dnmt3a*-mutant LSKs were biased toward early apoptotic states in MS5-Lx8 co-culture conditions (Figure 3E-G, Figure S4D). This differential pattern suggests that WT LSKs are more susceptible to progression toward irreversible apoptotic states in response to retrotransposon-induced inflammatory cues. Additionally, MS5-Lx8 exposure led to accumulation of WT LSKs in G1 phase and reduced progression into S-G2/M (Figure 3H-I, Figure S4E), indicating cell cycle arrest. In contrast, *Dnmt3a*-mutant LSKs displayed stable progression through the cell cycle with MS5-Lx8 exposure.

To minimize confounding effects from proliferative stress associated with *ex vivo* culture and more accurately assess HSC quiescence under inflammatory conditions, we evaluated the *in vivo* cell cycle response of WT and *Dnmt3a*-mutant HSCs following 7 days of recombinant IFNα or vehicle treatment. Notably, *Dnmt3a*-mutant HSCs did not alter their G0, G1, or S-G2/M states while WT HSCs exited G0 and accumulated in G1 phase following IFNα treatment (Figure 3J-K). Together, these studies demonstrate that *Dnmt3a*-mutant HSCs gain a competitive advantage over WT HSCs under retrotransposon-activated conditions by resisting alterations to their cell cycle and maintaining quiescence-enforcing programs following interferon stimulation.

### Retrotransposon-induced *Dnmt3a*-mutant HSC expansion is mediated through a viral mimicry response

While retrotransposons are closely linked to the activation of cGAS-STING and the RIG-I-like receptor pathways, whether retrotransposons promote CH through viral mimicry remains untested (Figure 4A). Thus, we sought to define the molecular pathways through which retrotransposons promote *Dnmt3a*-mediated clonal expansion. First, we confirmed that retrotransposons can broadly activate an IFN-I response across all major retrotransposon families in MS5 cells. In addition to a LINE-1 element (Lx8), we ectopically expressed a SINE element (B1) and an endogenous retrovirus element (MERV-L) that were similarly upregulated with aging in MSCs (Figure S1G, S4B, S5A). Indeed, all three retrotransposons increased *Ifnb1* mRNA expression and elevated IFNβ cytokine secretion (Figure S5B-C), confirming induction of an IFN-I response in MS5 cells. To functionally validate that IFN-I is the primary inflammatory cue promoting *Dnmt3a*-mutant cell expansion, we verified that there was no comparative induction of other CH- and aging-related inflammatory genes, including *Ifng, Il1b, Tnf*, and *Il6*, indicating that retrotransposons trigger inflammation in an IFN-I-specific manner (Figure S5D). Next, we tested whether treatment of MS5 cells with repeated doses of recombinant IFNα was sufficient to promote the fitness advantage of *Dnmt3a*-mutant cell *ex vivo* (Figure 4B). Indeed, sustained IFNα treatment of MS5 cells led to selective outgrowth of *Dnmt3a*-mutant LSK cells due to selective depletion of WT LSK cells (Figure 4C-D), phenocopying the effects of MS5-Lx8 co-culture (Figure 3C-D). These results show that retrotransposons act through chronic IFN-I to promote mutant HSC growth.

**Fig. 4.**
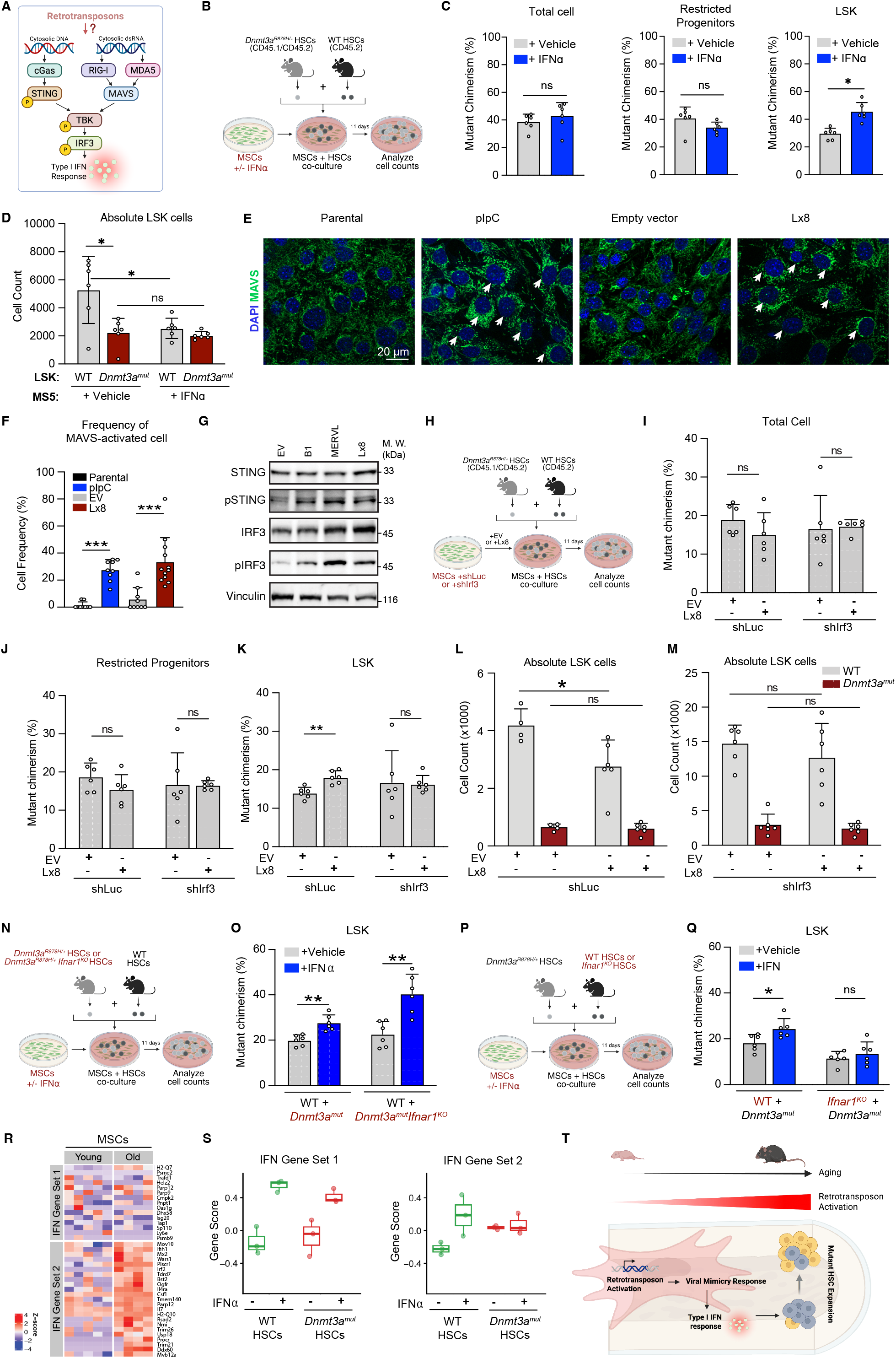
Retrotransposon-induced *Dnmt3a*-mutant HSC expansion is mediated through a viral mimicry response. **(A)** Pathway of retrotransposon-mediated viral mimicry response **(B)** Schematic of the *ex vivo* competitive co-culture model with 5ng/mL recombinant IFNα or vehicle treatment. LT-HSCs (CD150^+^CD34^−^LSKs) were sorted from WT CD45.2 mice and *Vav-iCre*;*Dnmt3a*^*fl-R878H*/+^ CD45.1/CD45.2 mice. WT and *Dnmt3a*-mutant LT-HSCs were mixed at a 2:1 ratio. After 11 days of co-culture, HSCs were analyzed by flow cytometry. **(C)** Flow cytometry analysis of mutant chimerism in total cells, restricted progenitors (LKs), and LSKs after *ex vivo* co-culture with or without 5ng/mL recombinant IFNα treatment (n=5-6). Mutant chimerism was calculated as the ratio of *Dnmt3a*-mutant CD45.1/CD45.2 cells to total cells within population. **(D)** Quantification of absolute LSK counts for mixed HSCs co-cultured with MS5 cells with or without 5ng/mL recombinant IFNα (n=5-6). Statistical significance for (C) and (D) was determined by multiple t-tests. **(E)** Immunofluorescent imaging of MAVS aggregation in parental MS5 cells and MS5 cells transfected with pIpC (positive control), EV, or Lx8. DAPI stained in blue. MAVS stained in green. White arrows identifying MAVS aggregated cells. **(F)** Quantification of the frequency of cells containing cytoplasmic MAVS aggregation. Statistical significance was determined by multiple t-tests. **(G)** Western blot analysis of STING and IRF3 phosphorylation in MS5 cells transfected with EV or retrotransposon constructs. **(H)** Schematic of the *ex vivo* competitive co-culture model using MS5 cells expressing shLuciferase (shLuc) control or shIrf3. **(I-K)** Flow cytometry analysis of mutant chimerism in total cells (I), restricted progenitors (J), or LSK cells (K) after *ex vivo* co-culture with MS5 cells expressing shLuciferase (shLuc) control or shIrf3 (n=5-6). Mutant chimerism was calculated as the ratio of *Dnmt3a*-mutant CD45.1/CD45.2 cells to total cells within population. Statistical significance was determined by multiple t-tests. **(L-M)** Quantification of absolute LSK counts for mixed HSCs co-cultured with MS5-EV or MS5-Lx8 cells expressing shLuc control (L) or shSting (M). Figures (A), (B), and (H) created with the aid of Biorender.

To further explore whether retrotransposon-induced inflammation is mediated by a viral mimicry response in MSCs, we directly interrogated the cGAS-STING pathway, which detects cytosolic dsDNA, and the RIG-I pathway, which detects cytosolic dsRNA (Figure 4A).^33-35^ Given that some cells silence these pathways, we first verified the intact nature of the cGAS-STING and RIG-I pathways in MS5 cells using dsDNA (via salmon sperm) or synthetic dsRNA (via polyinosine-polycytidylic acid), respectively (Figure S5E). Next, we found that retrotransposon upregulation not only leads to the cytoplasmic aggregation of MAVS, a mitochondrial adaptor protein that relays the signals from the RIG-I and MDA5 receptor activation, but it also leads to the phosphorylation of STING, the central mediator of the cGAS-STING pathway (Figure 4E-G). Hence, stromal cells can recognize and respond to multiple nucleic acid intermediates generated during the retrotransposon life cycle. Additionally, retrotransposon upregulation in MS5 cells induced phosphorylation of TBK1 and IRF3, which are both downstream mediators of the viral mimicry pathways (Figure 4G, Figure S5F).

To assess whether the viral mimicry response is necessary for retrotransposons to promote *Dnmt3a*-mutant expansion, we generated MS5 cells expressing either control shRNA (shLuciferase) or shRNA targeting *Irf3*. We validated that *Irf3* knockdown suppressed the IFN-I response (Figure S5G). We then overexpressed either Lx8 or EV control in these cells and performed competitive co-culture assays with a 1:2 mixed population of *Dnmt3a*-mutant and WT HSCs (Figure 4H). Ablation of *Irf3* in MS5 cells abrogated the *Dnmt3a-*mutant LSK expansion mediated by Lx8 overexpression, confirming that a viral mimicry response is required for retrotransposon-induced mutant LSK outgrowth (Figure 4I-K). Accordingly, the absolute number of *Dnmt3a*-mutant LSKs remained unchanged when co-cultured with MS5-EV or MS5-Lx8 following *Irf3* knockdown in the stromal cells (Figure 4L-M). Additionally, ablation of STING in MS5 cells similarly reduced the Lx8-mediated clonal advantage (Figure S5H-K). These results demonstrate that stromal expression of retrotransposons activate a cell-extrinsic viral mimicry response that promotes *Dnmt3a*-mutant HSC expansion.

### *Dnmt3a*-mutant HSCs selectively attenuate stromal IFN-I responses to gain a competitive advantage

Given that *Dnmt3a-*mutant HSCs demonstrate increased resistance to the inflammatory stress mediated by the viral mimicry response, we hypothesized that these cells may be functionally less responsive to stromal IFN-I signaling. To test this, we examined the effect of deleting the IFNα receptor in *Dnmt3a-*mutant cells (*Vav-iCre*;*Dnmt3*^*fl-R878Hl+*^;*Ifnar1*^*fl/fl*^) (Figure 5A). Strikingly, mutant HSCs lacking *Ifnar1* retained their growth advantage over WT HSCs under conditions of enhanced IFN-I stress (Figure 5B). In contrast, *Dnmt3a-*mutant HSCs with intact *Ifnar1* demonstrated no growth advantage over *Ifnar1-* deficient WT HSCs (*Vav-iCre*;*Ifnar1*^*fl/fl*^) under the same stress conditions (Figure 5C-D). We additionally confirmed that loss of *Ifnar1* itself did not alter the hematopoietic output of *Dnmt3a-*mutant HSCs (Figure S6A). Thus, WT HSCs are selectively disadvantaged by their intrinsic response to stromal IFNα signaling, whereas mutant HSCs gain a competitive advantage by attenuating their response.

**Fig. 5.** Wild-type HSCs have maladapted response to retrotransposon-mediated inflammation. **(A)** Schematic of the *ex vivo* competitive co-culture model in which WT HSCs are competed with *Dnmt3a*-mutant HSCs containing either an intact or deleted interferon alpha receptor (*Ifnar1)*. MS5 cells are treated with vehicle or 5ng/mL recombinant IFNα. **(B)** Flow cytometry analysis of mutant chimerism in LSKs after *ex vivo* co-culture as described in (A). **(C)** Schematic of the *ex vivo* competitive co-culture model in which *Dnmt3a*-mutant HSCs are competed either with WT HSCs or HSCs containing a deleted interferon alpha receptor (*Ifnar1)*. MS5 cells are treated with vehicle or 5ng/mL recombinant IFNα. **(D)** Flow cytometry analysis of mutant chimerism in LSKs after *ex vivo* co-culture as described in (C). **(E)** Heatmap of interferon alpha response genes in young and old MSCs and HSCs. **(F)** Differential activation of IFN gene sets defined in (E) for WT and *Dnmt3a*-mutant HSCs following IFNα or vehicle treatment. **(G)** Gene expression of *Ifnar1* and *Ifnar2* derived from normalized counts using RNA-seq data of WT and *Dnmt3a*-mutant HSCs following IFNα or vehicle treatment. **(H)** Proposed model for retrotransposon-driven clonal hematopoiesis during aging. Statistical significance for (B) and (D) was determined by multiple t-tests. Figures (A), (C), and (H) created with the aid of Biorender.

### Wild-type HSCs have maladapted response to retrotransposon-mediated inflammation

To dissect the intrinsic differences in IFN-I responsiveness, we leveraged our bulk RNA sequencing data of young and aged bone marrow cell types (Figure 1A) to identify subsets of IFN-I genes that were either associated with the aged MSC or the aged HSPC interferon program (Figure 5E). We then performed bulk RNA sequencing on WT and *Dnmt3a-*mutant HSCs exposed to IFNα or vehicle. Following IFNα treatment, WT HSCs upregulated IFN gene sets associated with both aged MSCs and HSPCs. In contrast, *Dnmt3a-*mutant HSCs appeared to only upregulate a smaller subset of IFN genes most closely associated with MSC aging (Figure 5F). WT and *Dnmt3a-*mutant HSCs expressed the same levels of *Ifnar1* and *Ifnar2*, ruling out receptor-level differences in IFN-I responsiveness (Figure 5G). Thus, we propose that mutant HSCs are not globally insensitive to IFN-I signaling but rather are selectively unresponsive to a specific interferon program.

In summary, our data support a model in which aging upregulates retrotransposons in MSCs, triggering an IFN-I response via viral mimicry that selectively suppresses WT HSC fitness while sparing *Dnmt3a*-mutant HSCs, thereby promoting CH (Figure 5H).

## Discussion

Our work identifies a previously unrecognized role for retrotransposons in shaping HSC clonal dynamics through MSCs, highlighting a non-cell-autonomous mechanism by which the aged niche promotes CH. We show that aging induces a retrotransposon-mediated IFN-I response that selectively impairs WT HSCs while permitting *Dnmt3a*-mutant HSCs to maintain growth and self-renewal. This aligns with previous work establishing that chronic IFN-I signaling promotes HSC exhaustion, leading to loss of the HSC pool.^36-38^ This distinction between mutant and WT HSC response emphasizes how clonal dominance can emerge from differential resilience to environmental stress in conjunction with inherent hyperproliferative properties of mutant clones. Our work also highlights differences between the role of type I and type II interferons in clonal hematopoiesis: In contrast to IFNγ, which directly promotes Dnmt3a-mutant expansion, our data indicate that IFNα signaling within mutant HSCs is largely dispensable.^3,39^ Indeed, we show that *Dnmt3a*-mutant HSCs are more resistant to IFN-I-induced cell cycle disruptions and retain quiescence in the face of inflammatory challenge. Importantly, Dnmt3a-mutant HSCs are not globally insensitive to IFN-I but instead selectively evade specific interferon programs, allowing them to remain responsive to some cues while avoiding those that suppress WT HSC fitness. Together, this supports a growing model in which resistance to inflammation is a key determinant of mutant clonal advantage.^5,13,37,39,40^ Mechanistically, we also show that retrotransposon-induced inflammation in the aged bone marrow is driven by stromal viral mimicry. While viral mimicry is well known as a *cell-intrinsic* driver of interferon signaling, our study reveals a distinct, *cell-extrinsic* role in promoting somatic selection.^19,21^

These findings have several implications. Targeting retrotransposon activity or downstream pathways (e.g., reverse transcriptase inhibitors or cGAS–STING blockade) may mitigate age-associated CH. Second, our work provides key evidence that *Dnmt3a*-mutant cells are resistant to IFN-I signaling, inviting future studies to explore the mechanisms of such resistance. Thirdly, while our study focused on *Dnmt3a*-mutant HSCs, future studies should test whether HSCs harboring *TET2, ASXL1*, or *TP53* mutations similarly benefit from retrotransposon-driven inflammation, and whether they engage distinct or overlapping pathways.

In conclusion, our study identifies retrotransposon activation in the aged bone marrow stroma as a key extrinsic force driving CH through viral mimicry pathways. These findings highlight the importance of the aging bone marrow environment in shaping hematopoietic evolution and provide a mechanistic framework linking retrotransposons, innate immunity, and somatic selection in aging.

## Supporting information

SuppFigs

## Acknowledgements

We thank all the members of the Xu lab for critical discussions, contributions, and support. We thank the Children’s Research Institute at UT Southwestern for use of the Moody Flow Cytometry Facility and Sequencing Core. We thank Andrew Koh (UTSW), Stephen Skapek (Duke), and Stephen Chung (UTSW) for helpful discussions and support. This work was supported by funding provided by Hyundai Hope on Wheels, Children’s Cancer Fund, and CPRIT (RP220375).

## Author Contributions

**Q.Z**.: investigation, methodology, analysis, writing; **W.H**.: investigation, analysis; **J.T**.: investigation, analysis: **M.L**.: analysis; **S.R**.: investigation; **Y.J.K:** investigation; **D.C**.: investigation; **K.D**.: editing; **J.T**.: methodology; **J.X**.: conceptualization, methodology, editing; **S.Z**.: conceptualization, investigation, methodology, analysis, writing

