## Supplementary material for "Retrotransposons Promote *Dnmt3a*-Mutant Clonal Hematopoiesis Through Aging-Related Stromal Inflammation": SuppFigs

**FIGURE S1.**

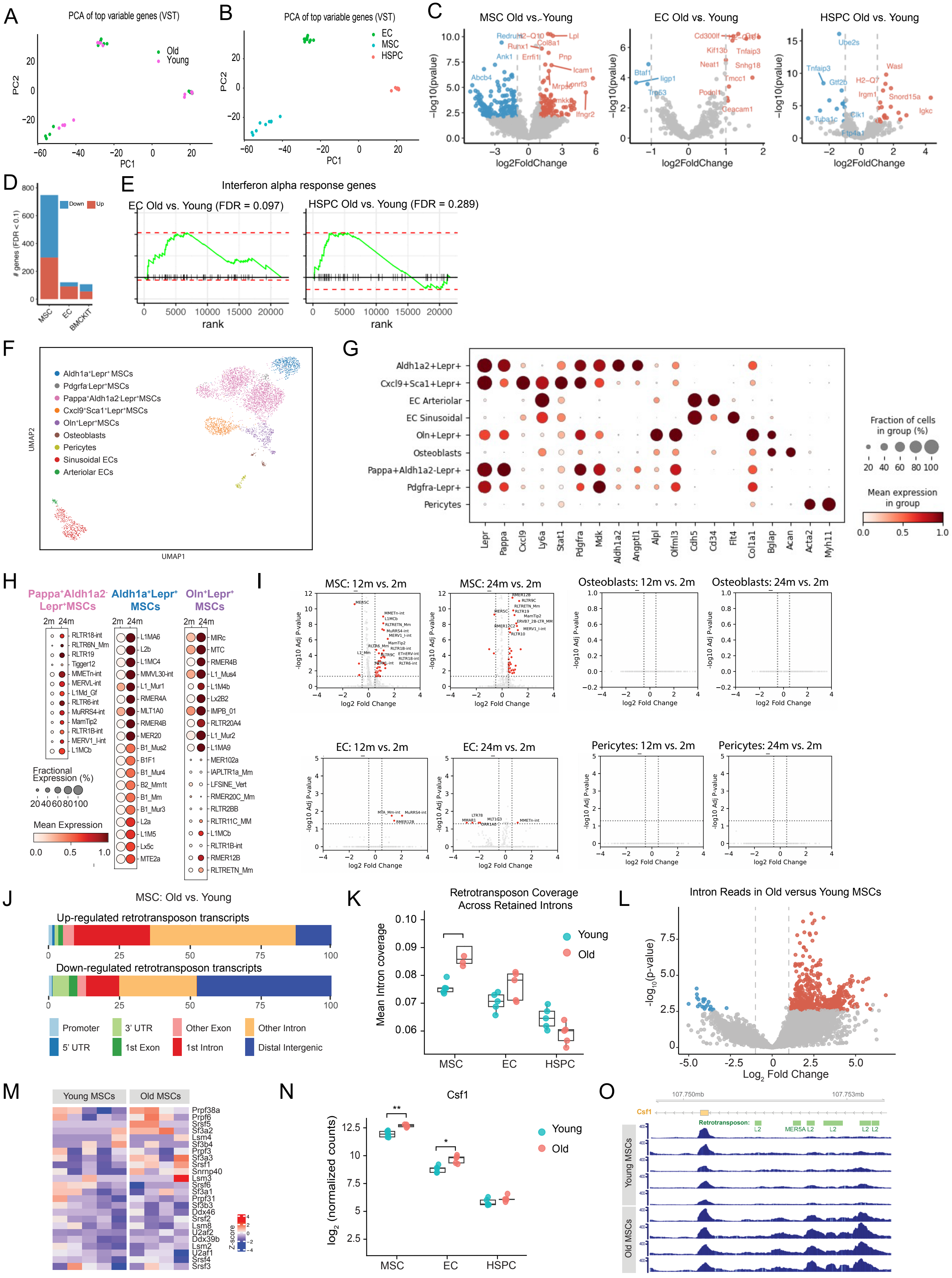

A
PCA of top variable genes (VST)
Old
Young
PC1
PC2
B
PCA of top variable genes (VST)
EC
MSC
HSPC
PC1
PC2
C
MSC Old vs. Young
EC Old vs. Young
HSPC Old vs. Young
-log10(pvalue)
log2FoldChange
D
Down
Up
### genes (FDR < 0.1)
MSC
EC
BMCKIT
E
Interferon alpha response genes
EC Old vs. Young (FDR = 0.097)
HSPC Old vs. Young (FDR = 0.289)
rank
F
Aldh1a⁺Lepr⁺ MSCs
Pdgfra⁻Lepr⁺MSCs
Pappa⁺Aldh1a2⁻Lepr⁺MSCs
Cxcl9⁺Sca1⁺Lepr⁺MSCs
Oln⁺Lepr⁺MSCs
Osteoblasts
Pericytes
Sinusoidal ECs
Arteriolar ECs
UMAP1
UMAP2
G
Aldh1a2+Lepr+
Cxcl9+Sca1+Lepr+
EC Arteriolar
EC Sinusoidal
Oln+Lepr+
Osteoblasts
Pappa+Aldh1a2-Lepr+
Pdgfra-Lepr+
Pericytes
Lepr
Pappa
Cxcl9
Ly6a
Stat1
Pdgfra
Mdk
Aldh1a2
Angptl1
Alpl
Olfml3
Cdh5
Cd34
Flt4
Col1a1
Bglap
Acan
Acta2
Myh11
Fraction of cells in group (%)
Mean expression in group
H
Pappa⁺Aldh1a2⁻Lepr⁺MSCs
Aldh1a⁺Lepr⁺ MSCs
Oln⁺Lepr⁺ MSCs
2m 24m
Fractional Expression (%)
Mean Expression
I
MSC: 12m vs. 2m
MSC: 24m vs. 2m
Osteoblasts: 12m vs. 2m
Osteoblasts: 24m vs. 2m
EC: 12m vs. 2m
EC: 24m vs. 2m
Pericytes: 12m vs. 2m
Pericytes: 24m vs. 2m
-log10 Adj P-value
log2 Fold Change
J
MSC: Old vs. Young
Up-regulated retrotransposon transcripts
Down-regulated retrotransposon transcripts
Promoter
5' UTR
3' UTR
1st Exon
Other Exon
1st Intron
Other Intron
Distal Intergenic
K
Retrotransposon Coverage Across Retained Introns
Mean Intron coverage
Young
Old
MSC
EC
HSPC
L
Intron Reads in Old versus Young MSCs
-log10(p-value)
Log2 Fold Change
M
Young MSCs
Old MSCs
Z-score
N
Csf1
log2 (normalized counts)
Young
Old
MSC
EC
HSPC
O
107.750mb
107.753mb
Csf1
Retrotransposon:
Young MSCs
Old MSCs

**FIGURE S2.**

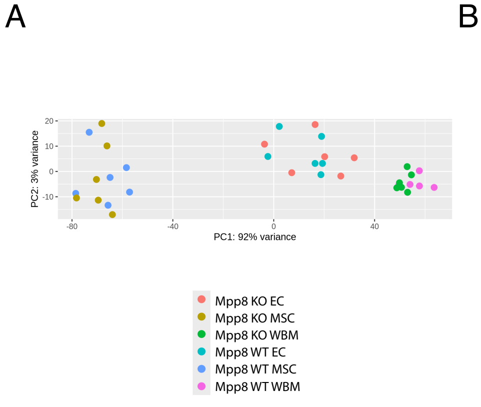


B

| Gene | log2FoldChange | pvalue |
|---|---|---|
| L1Lx_III:L1:LINE | 4.51191295 | 0.00421304 |
| RLTR6B_Mm:ERV1:LTR | 3.8593653 | 0.00099708 |
| Lx7:L1:LINE | 3.60118742 | 0.00516638 |
| L1_Mur2:L1:LINE | 3.44331312 | 0.00308796 |
| MT-int:ERVL-MaLR:LTR | 3.2970531 | 0.01030051 |
| L1MdV_I:L1:LINE | 3.27881583 | 0.01284158 |
| Lx5c:L1:LINE | 3.1929477 | 0.04530335 |
| RMER3D-int:ERVK:LTR | 3.05251821 | 0.03630125 |
| RLTR33:ERVK:LTR | 2.91523052 | 0.01455369 |
| Lx5b:L1:LINE | 2.91333384 | 0.00207954 |
| L1MdA_VII:L1:LINE | 2.90651458 | 0.00559047 |
| L1MdA_VI:L1:LINE | 2.65235907 | 0.00251206 |
| L1MdFanc_I:L1:LINE | 2.50524765 | 0.00717414 |
| GSAT_MM:Satellite:Satellite | 2.45419879 | 0.00085326 |
| B1_Mm:Alu:SINE | 2.42847489 | 0.00040799 |
| L1MdA_V:L1:LINE | 2.31766455 | 0.02974697 |
| Lx5:L1:LINE | 2.09693494 | 0.01257358 |
| L1MdMus_I:L1:LINE | 1.9360755 | 0.01089427 |
| L1MdV_II:L1:LINE | 1.86060853 | 0.00205859 |
| L1MdGf_II:L1:LINE | 1.70011986 | 0.00125517 |
| L1MdF_III:L1:LINE | 1.66969552 | 0.00722923 |
| L1Lx_I:L1:LINE | 1.32224335 | 0.03317401 |
| L1MdMus_II:L1:LINE | 1.30049941 | 0.04248491 |
| B1_Mur3:Alu:SINE | 1.24364665 | 0.02342118 |
| L1MdA_III:L1:LINE | 1.15383113 | 0.00942794 |
| B1_Mus1:Alu:SINE | 1.14958765 | 0.040341 |
| RLTR10:ERVK:LTR | 1.00291797 | 0.05145817 |
| L1MdF_I:L1:LINE | 0.83976363 | 0.04611251 |
| L1MdTf_I:L1:LINE | 0.83407284 | 0.0385233 |
| RLTR4_MM-int:LTR:LTR | 0.71797484 | 0.00287472 |
| L1MdTf_III:L1:LINE | 0.51597094 | 0.04046438 |
| L1MdA_I:L1:LINE | 0.51046779 | 0.04618635 |

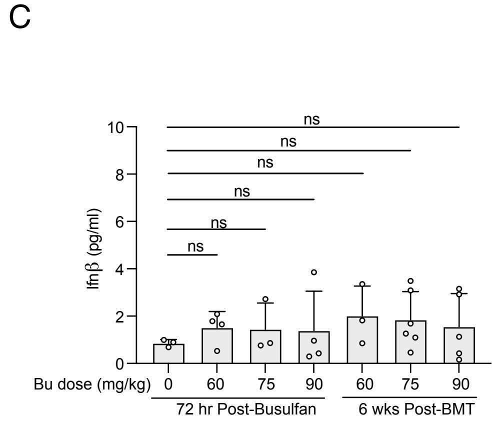


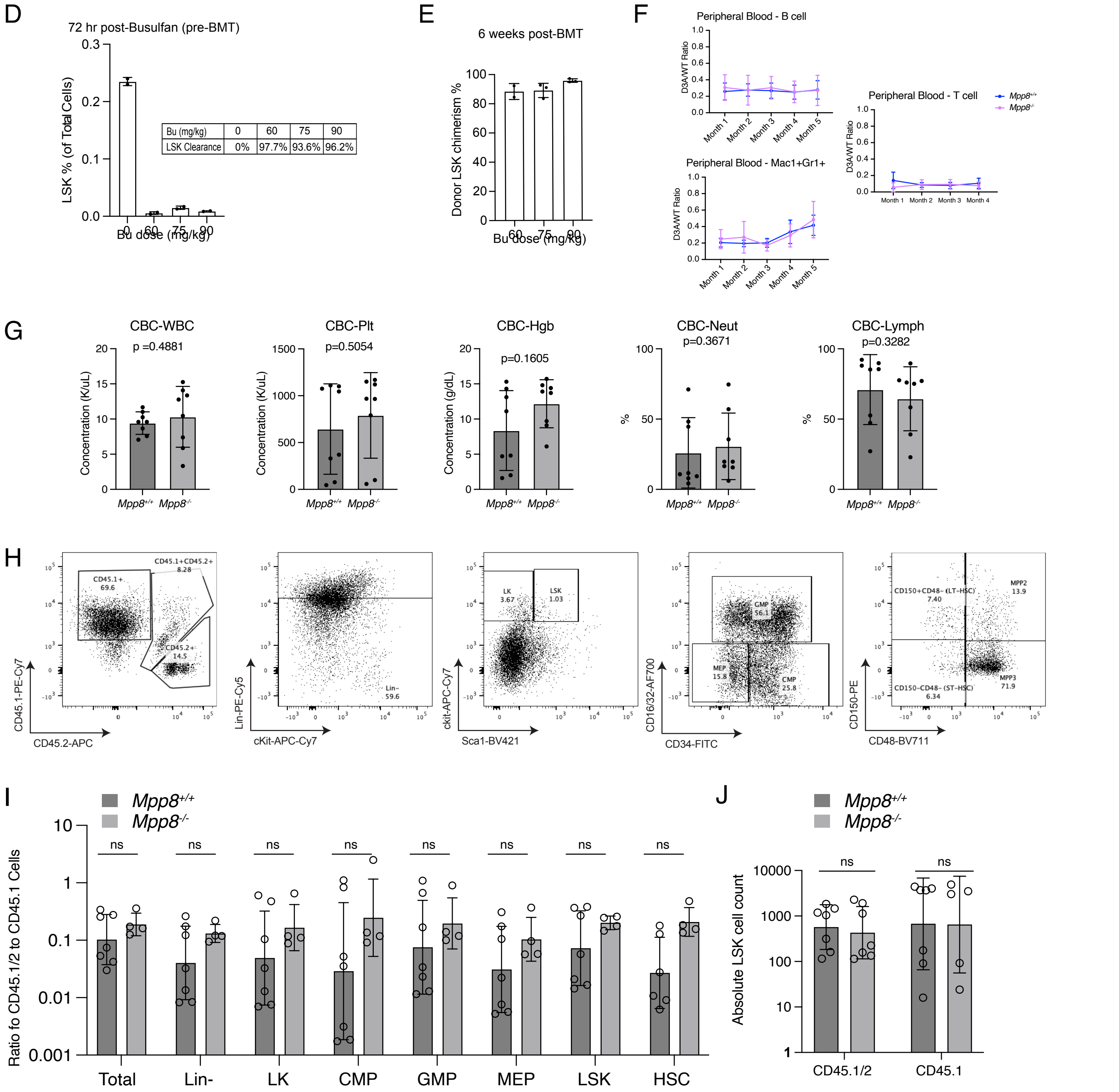

**FIGURE S3.**

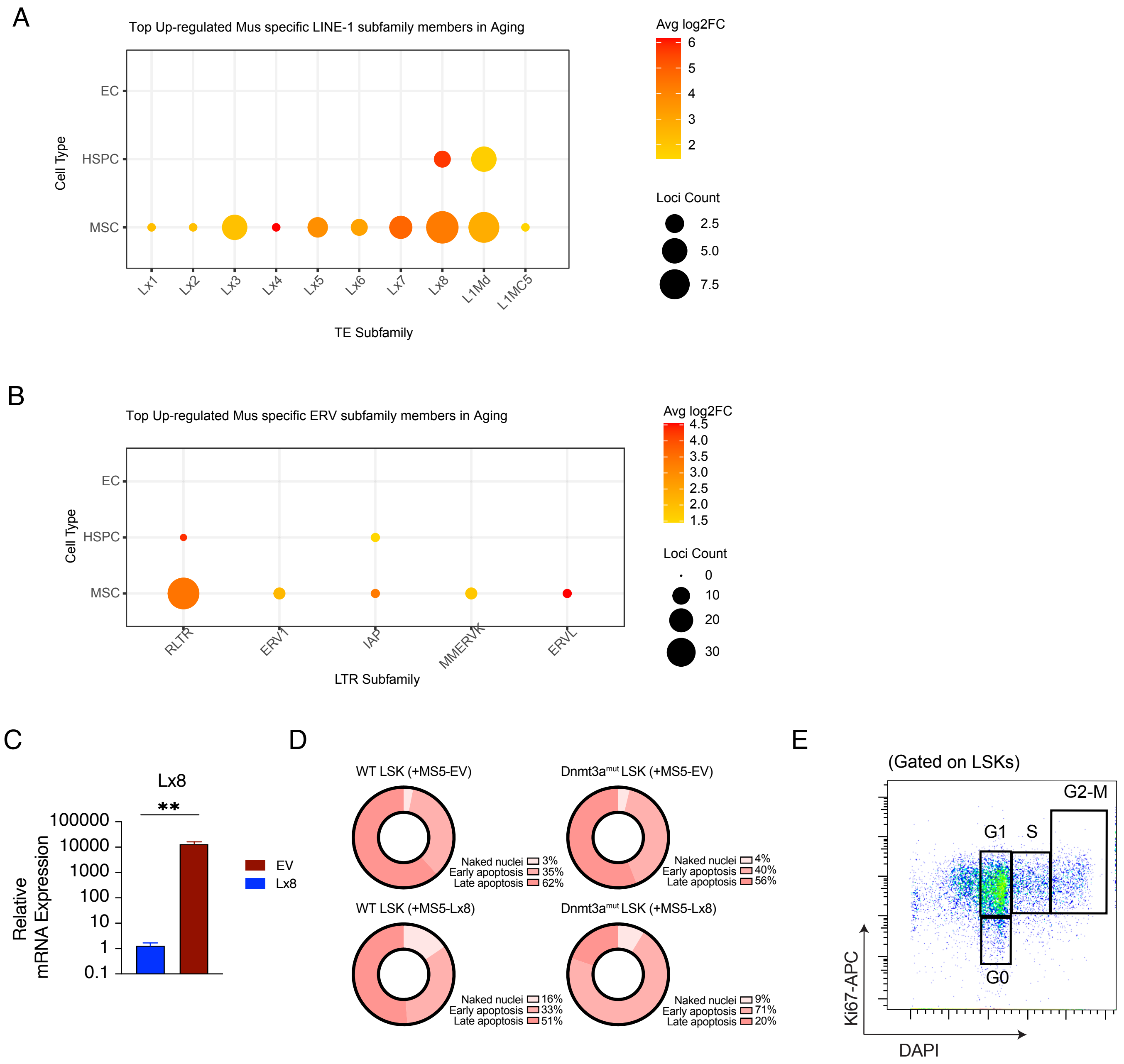

A
Top Up-regulated Mus specific LINE-1 subfamily members in Aging
Cell Type
EC
HSPC
MSC
Lx1
Lx2
Lx3
Lx4
Lx5
Lx6
Lx7
Lx8
L1Md
L1MC5
TE Subfamily
Avg log2FC
6
5
4
3
2
Loci Count
2.5
5.0
7.5
B
Top Up-regulated Mus specific ERV subfamily members in Aging
Cell Type
EC
HSPC
MSC
RLTR
ERV1
IAP
MMERVK
ERVL
LTR Subfamily
Avg log2FC
4.5
4.0
3.5
3.0
2.5
2.0
1.5
Loci Count
0
10
20
30
C
Lx8
**
100000
10000
1000
100
10
1
0.1
Relative mRNA Expression
EV
Lx8
D
WT LSK (+MS5-EV)
Naked nuclei 3%
Early apoptosis 35%
Late apoptosis 62%
Dnmt3a^mut LSK (+MS5-EV)
Naked nuclei 4%
Early apoptosis 40%
Late apoptosis 56%
WT LSK (+MS5-Lx8)
Naked nuclei 16%
Early apoptosis 33%
Late apoptosis 51%
Dnmt3a^mut LSK (+MS5-Lx8)
Naked nuclei 9%
Early apoptosis 71%
Late apoptosis 20%
E
(Gated on LSKs)
G2-M
G1
S
G0
Ki67-APC
DAPI

**FIGURE S4.**

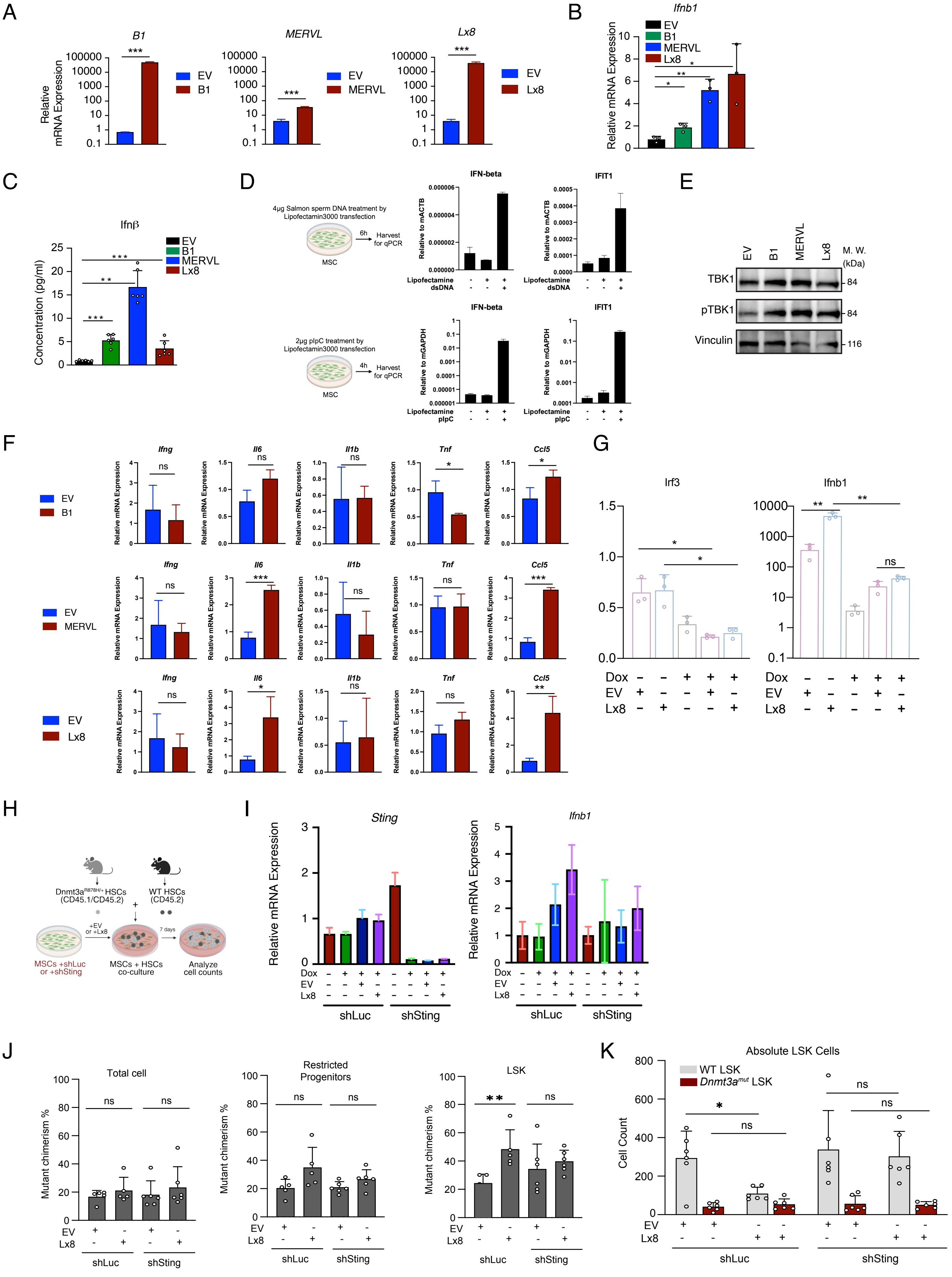

A
B1
MERVL
Lx8
Relative mRNA Expression
EV
B1
MERVL
Lx8
***
B
Ifnb1
Relative mRNA Expression
C
Ifnβ
Concentration (pg/ml)
D
4μg Salmon sperm DNA treatment by Lipofectamin3000 transfection
6h
Harvest for qPCR
MSC
IFN-beta
IFIT1
Relative to mACTB
Lipofectamine
dsDNA
2μg pIpC treatment by Lipofectamin3000 transfection
4h
Relative to mGAPDH
pIpC
E
TBK1
pTBK1
Vinculin
M. W. (kDa)
84
116
F
Ifng
Il6
Il1b
Tnf
Ccl5
ns
G
Irf3
Ifnb1
Dox
EV
Lx8
H
Dnmt3a^R878H/+ HSCs (CD45.1/CD45.2)
WT HSCs (CD45.2)
+EV or +Lx8
7 days
MSCs +shLuc or +shSting
MSCs + HSCs co-culture
Analyze cell counts
I
Sting
shLuc
shSting
J
Total cell
Restricted Progenitors
LSK
Mutant chimerism %
K
Absolute LSK Cells
WT LSK
Dnmt3a^mut LSK
Cell Count
